# Asynchronous population dynamics induced by higher-order and negative asymmetric ecological interactions

**DOI:** 10.1101/2025.08.21.671436

**Authors:** Dweepabiswa Bagchi, P. K. Nuha Fatima

## Abstract

Phase synchronized population dynamics of various species constituting a complex ecosystem elevates the risk of their extinction due to both environmental stochasticity and simulateneous low density fluctuations. Therefore, an extremely vital approach to measure the extinction risk of an ecosystem as a whole is to quantify the phase synchrony among the species populations co-habiting and interacting with each other in an ecosystem. Generally, in models describing population dynamics of ecosystems, both trophic and non-trophic inter-species interactions are modelled as interactions between two species. This approach contradicts the fact with such a large number of species living in close proximity, more than two species must partake in the same interaction influencing the population dynamics of each other. To address this, higher-order interactions need to be incorporated in the models describing population dynamics of an ecosystem. Consequently, their effect on phase synchronization of populations also need to be investigated. In this study, we model a species-rich ecosystem as a complex phase oscillator network and examine the phase dynamics of the total population. Each node of this network represents a constituent species, modelled as a Sakugachi-Kuramoto phase oscillator coupled non-linearly to the other nodes through both first-order and higher-order inter-species interactions. These interactions can be both mutualistic (positive) and antagonistic (negative) in nature. Along with the higher-order interactions, we also incorporate inherent asymmetry among the nodes to account for habitat heterogeneity. Further, we investigate the effects of both higher-order coupling and asymmetry on the phase synchronization of the total population. Our findings demonstrate that higher-order interactions above a threshold amplitude enforces a transition from synchronous to asynchronous dynamics of the ecosystem. Further, we find that increase in the size and diversity of the ecosystem leads to an increase in the threshold value of higher order coupling required to reach asynchronous dynamics. We also demonstrate that this higher-order induced asynchrony is further promoted by high asymmetry among the individual nodes. Importantly, negative inter-species interactions, if existing to a high degree also induce asynchrony in the system. Moreover, the size of the network also plays a role in deciding the threshold value of higher order coupling required to induce asynchrony.

## I. INTRODUCTION

Ecosystems consist of populations of multiple interacting species living in close proximity. A paradigm of modelling these populations are prey-predator models[1, 2]. Often in prey-predator models the prey and predator populations exhibit oscillatory dynamics and behave as a single oscillator [3]. For a spatially extended ecosystem, such oscillators are coupled with one another through processes like dispersal [4], cross-trophic interactions or other competitive and/or mutualistic interactions [6, 7]. Ecosystems are thus accurately described as complex networks formed by coupled oscillators [8], where the total population dynamics of the network is governed by non-linear, coupled differential equations. As species-richness (i.e., number of species) in an ecosystem increases, the number of coupled non-linear equations increase and the complex parameterisation and limited analytical scope of these equations pose a serious challenge to accuracy of predictions. To address this, modelling of population of large ecological networks formed by various species needs to be simplified without losing the essential dynamical features of these models, and specific focus need to be maintained on measures quantifying extinction risk of a large population forming a large ecological network. It is well established that when all species residing in an ecosystem have phase synchronized population dynamics, the persistence of the ecosystem is lowered, while asynchronous population dynamics of the constituent species denote greater resilience of an ecosystem[9, 10]. Therefore, an essential approach to quantify the extinction risk of the population of a large ecological network is to investigate the phase synchronization of the total population dynamics of the network. For investigating phase phase dynamics of large populations of coupled oscillators, the Sakaguchi-Kuramoto model, focusing on the phases of oscillators and the synchronisation of the phases, is an extremely powerful framework [11].

Recent studies have used the Sakaguchi-Kuramoto model, to illustrate significant ecological effects [12], and [6, 13] have demonstrated that there exists strong similarities in dynamical behavior of a population described by the paradigmatic prey-predator Rozensweig-McArthur model and the dynamics described by the Sakaguchi-Kuramoto model. Thus, prey-predator pairs can be effectively modelled using the Sakaguchi-Kuramoto framework, with special focus on the phase dynamics of the population. Vandermeer [13] has reported that weak cross-feeding leads to in-phase synchrony among the oscillators, while competitive interactions result in antiphase synchrony, both of which are types of phase synchronizations nevertheless. Empirical confirmation of oscillatory dynamics in such qualitative coupling arrangements has also been observed by Beninc’a et.al.[14], cementing the importance of the phase-oscillator approach in modelling populations of large ecological networks. Therefore, an ecosystem can be described as a large ecological network, which in turn consists of non-linearly coupled Sakaguchi-Kuramoto phase oscillators. In our study, we take this well-established approach. Another important result that has been reported by numerous studies is that close proximity of mutltiple species in a species-rich ecosystem ensures that various inter-species interactions are essentially not first-order in nature, but rather involve more than two species simultaneously [15– 18].

Motivated by the above, in this study we consider the inclusion of higher-order (second-order) inter-species interactions of both trophic and non-trophic nature. To account for the habitat heterogeneity found in a natural ecosystem, we incorporate an asymmetric phase [19] in the dynamics of the constituent oscillators or nodes forming the ecological network, i.e., we allow each of the oscillators or nodes to possess varying phases even as they tend to synchronize. We introduce an adjacency matrix is defined and introduced to detail the interactions (couplings) between the constituent nodes of the network. In this study, the interactions we consider between two species in are not exclusively trophic in nature. Other prevalent ecological interactions like cross-feeding, resource competition, mutualism, commensalism and others involving more than one pair of species [5, 6] are also incorporated into the model. Notably, these interactions either benifits each of the species involved mutually (i.e., postive) or antagonizes all the involved species (i.e., negative). Our study demonstrates that incorporating the aforementioned modifications to the phase oscillator model leads to overall asynchronous dynamics for the entire network. Asynchronous dynamics is associated directly with beneficial conditions within the ecosystem, often due to creation of a source-sink meta-community [4]. Therefore, we claim that the continued existence of numerous ecological networks may quite possibly be due to the overall network dynamics of an ecological network being asynchronous due to the influuence of higher-order interactions. Simultaneously, this result very strongly indicates the presence of large magnitude of higher-order interactions in real ecosystems, as was a major consideration of our modelling approach.

## II. MODELS

As discussed above, we construct an ecological network, where each node of the network behaves as a phase-oscillator governed by the higher order Sakaguchi-Kuramoto model. To account for multi-species interactions and the inbuilt heterogeneity of a spatially extended natural ecosystem, we introduce asymmetry and higher-order interactions. To describe ecological networks, we consider two kinds of topology. First, we consider globally coupled networks (i.e., all nodes are interconnected), and then we extend our results to consider a random network structure, created using Erdös–Rényi random network formalism with finite connectivity probability. Both these network structures are quantified by an adjacency matrix detailing the coupling between the oscillators. Factoring in all this, the combined phase dynamics of the total ecological network formed by coupled phase oscillators, is governed by the equation [20, 21],

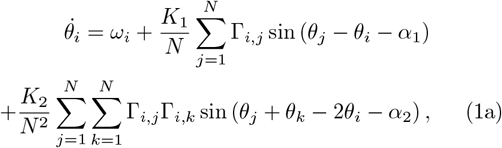

where, 0 ≤ *α* ≤ π/2 is the asymmetric phase lag parameter which introduces a phase delay among the interacting oscillators, *K*_1_ and *K*_2_ are the 1st and second-order coupling strength terms and the adjacency matrices *γ*_*i*_, *j* and *γ*_*i*_, *k* captures the interactions between oscillators *i,j* and *i,k* respectively. By adjusting the value of *α*_1_ and *α*_2_, we introduce varying degrees of asymmetry in the coupling. For this study, all *ω*_*i*_s were chosen at random from a normal distribution with standard deviation of 0.001. The initial conditions, for 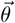 (i.e., phase values) were generated randomly. The degree of phase synchrony of the entire ecological phase-oscillator network is quantified by the order parameter measure given by,

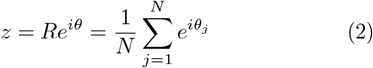

Here, *R* is the vector sum of the individual vectors and Θ is the average phase. When mod *z* = 1, the oscillators are fully phase synchronised and when mod *z* = 0, the oscillators are anti-phase synchronised or completely asynchronous. Throughout this work, *ω*_*i*_ is fixed at 1. Notably, we obtained the same results for any constant value of *ω*_*i*_.

### A. Positive and negative interactions

In the introduction, we highlighted the presence of various interactions within the same network. In the formalism we took from Fooroshani and Vandermeer [22], mutualistic interactions are considered positive interactions (+) which have a synchronizing effect between the nodes, while antagonistic interactions are considered negative interactions (-) have an asynchronous effect on the network dynamics.[23] states that stable equilibrium is maintained through positively coupld oscillators synchronizing with one another while avoiding proximity to negatively coupled counterparts. To account for such various possible inter-species interactions (couplings) naturally occuring in an ecosystem, we further incorporate both positive and negative interactions, manifested through both (+ve) and (-ve) *K*_1_ and *K*_2_ values in the network dynamics.

## III. RESULTS

### A. Effect of increasing asymmetry on the network dynamics

Figure. 1 demonstrates the effect of increasing first-order coupling (*K*_1_) on a network of 50 nodes. Fixing the first-order asymmetric (*α*_1_) phase parameter value at 0.1, we vary the first-order coupling *K*1 across the range 0 − 0.5. With increasing coupling strength, the network exhibits strong phase synchrony (*Z* ≈ 1). However, with increased asymmetry in phase (*α*_1_ = π/4 ≈ 0.78), the collective transition of oscillators to phase synchrony starts to dissipate. For this higher value of asymmetry, perfect phase synchronization is lost and partial phase synchronization is exhibited. Figure. 1(b) demonstrates that inclusion of only asymmetry in the dynamics can lower the degree of phase synchrony of the network but cannot lead to asynchronous dynamics. Next, we investigate the effect of increasing network size and varied network structure on this trend.

**FIG. 1.**
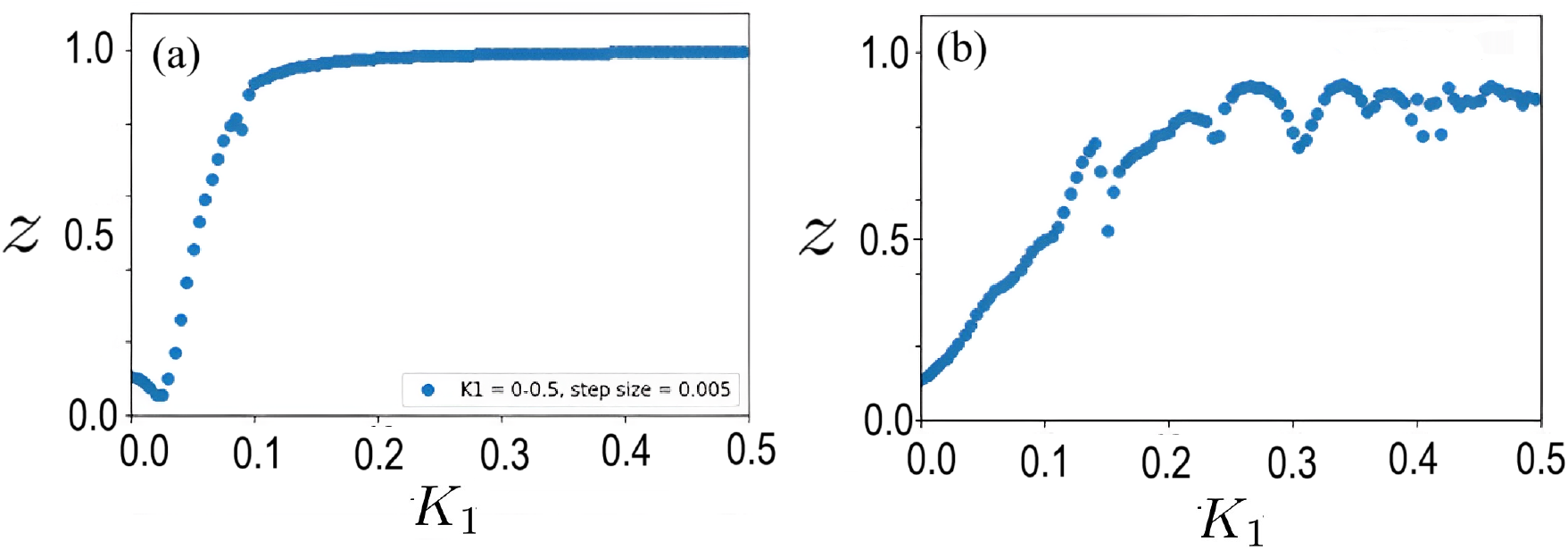
The effect of different values of first-order coupling strength (*K*_1_) on phase synchronization (*Z*) of the ecological network for a) low first order assymetry (*α*_1_ = 0.1), b) and high first-order assymetry ( *α* = *π/*4 ≈ 0.78). Higher *α*_1_ values break the symmetry of synchronisation

Figure. 2(a) demonstrates the behaviour of the phases of a globally coupled network of 20 nodes, for moderate first-order asymmetry (*α*_1_ = π/4 ≈ 0.78). When asymmetry is very high, the oscillators form 2 clusters. Even though the oscillators in a single cluster are phase synchronized (with phases varying over a limited range), the two clusters are individually anti-phase synchronized,lowering the phase synchorny of the dynamics. Notably, there are also few oscillators which do not cluster with any of the groups. Such elements are called chimeras. As demonstrated by Fig. 2(b), with increasing size of globally coupled networks, the distribution of both the phase synchronized cluster begins to increase. A much more prominent change in noticed in case of random networks. In case of high asymmetry, the clusters are numerous in number, smaller in size and in fact, slightly increases with increasing network size. In case of random networks, figures 2(c) and 2(d) demonstrate an intriguing relationship between high value of *α*_1_ and the synchronization patterns of the network, suggesting the existence of a critical threshold, after which higher values of *α*_1_ leads to a transition of the network dynamics from synchrony to complete asynchrony. Notably however, the asymmtery required for these networks to exhibit asynchronous dynamics is pretty high, and we will elaborate more on that in the following. Our results demonstrate the emergence of anti-phase group dynamics in the network when the network topology is global, and the emergence of chimeric elements and singular splay states when the network topology is represented by a random graph. Fig. 2 also showcases the distribution and arrangement of clusters and chimeric elements within the networks. The asynchrony is pronounced in case of random networ structure. Next, we introduce higher-order coupling into the system and analyze it’s effects on the phase synchrony of the entire network. We consider global networks primarily for observing the impacts of higher-order interactions/coupling on phase asynchrony of the system.

**FIG. 2.**
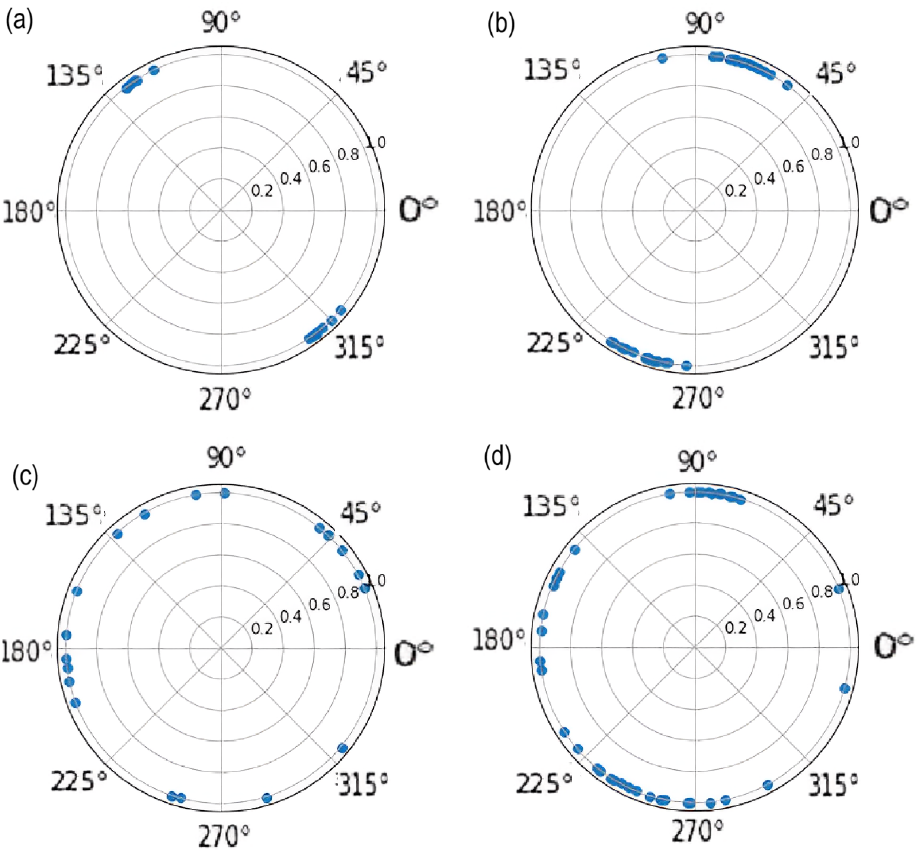
Clusters of phase synchronized oscillators, i.e., synchrony groups are formed for high asymmetry (*α*_1_ = *π/*4 ≈ 0.78) and existence of chimeric elements found for global network with (a) 20 and (b) 40 nodes. More prominent asynchronization, for same asymmetry is found for random network with (c) 20 and (d) 40 nodes. Asynchrony is more prominent in case of random network.

### B. Effect of higher-order coupling on network dynamics

As discussed in the introduction, to realistically model an ecological system it is imperative to consider more than 2 species as part of the same interaction. Accordingly, we introduce the higher-order (second-order) interaction between different oscillators on the dynamics of the ecological network with the amplitude of both first-order (*K*_1_) and higher-order coupling (*K*_2_) being comparable and investigate the impact of their interplay. The introduction of second-order interactions drastically disrupts the phase synchrony within the network, leading to overall asynchronous network dynamics when the coupling altitude reaches a value higher than a threshold. Fig. 3(a) demonstrates for a realistic value of *K*_1_, the decrease of order parameter *Z* of a globally coupled 20 node network with increasing higher order coupling. There is a transition of order parameter from high value (*Z* ≈ 1.0, signifying perfect phase synchrony between the nodes) to very low value (*Z* ≈ 0.2, signifying very low phase synchrony between the nodes). Eventually the order parameter *Z* approaches zero when the higher-order coupling increases beyond a threshold value. This also corresponds to the formation of distinct clusters (and chimeric elements) of phase-synchronized oscillators. However, their number are much smaller compared to the previous case where no higher-order coupling existed.

**FIG. 3.**
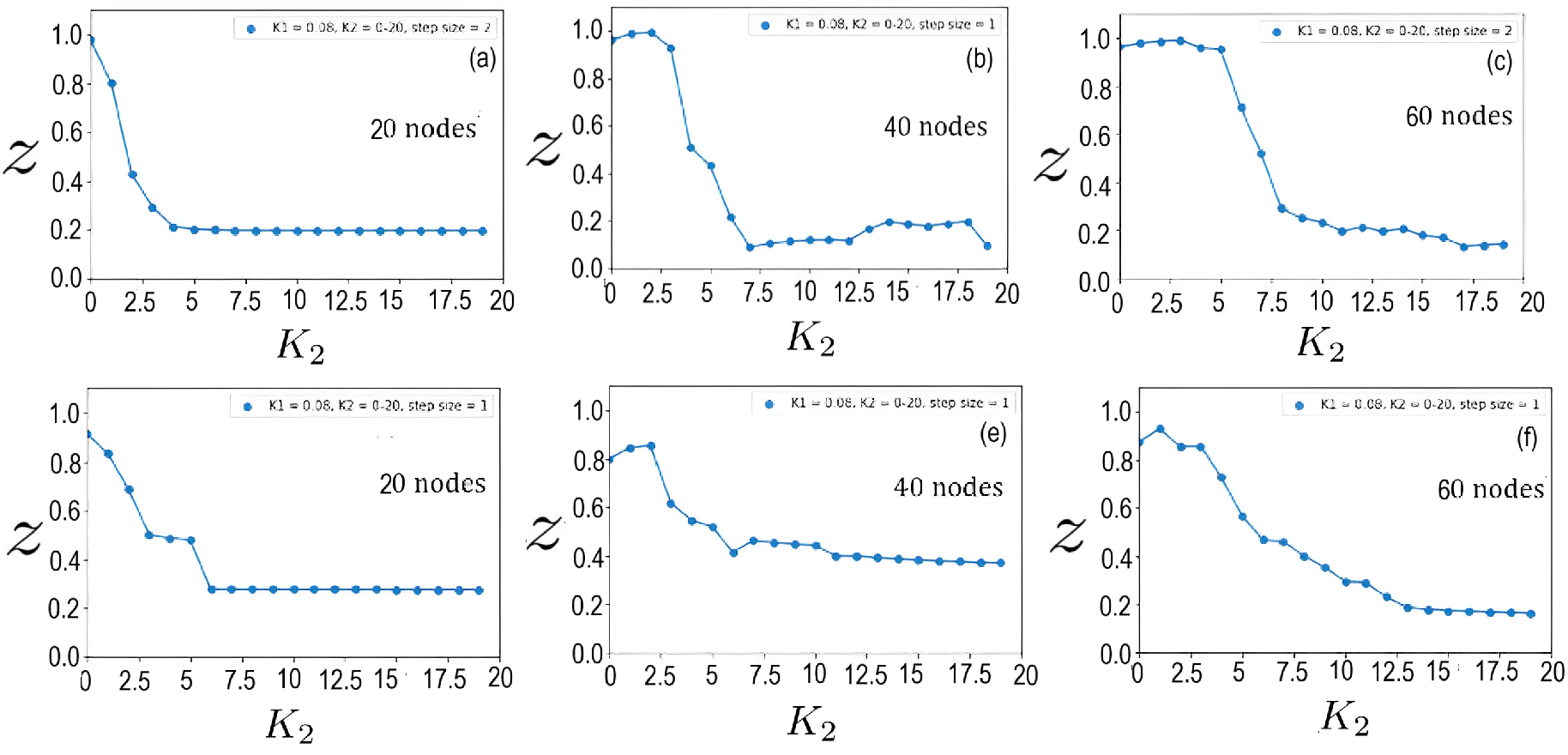
Order parameter (*Z*) vs higher-order coupling (*K*_2_) plot for global network of with (a) 20, (b) 40 and (c) 60 nodes. The same is done for random network with (d) 20, (e) 40 and (f) 60 nodes. The parameter values are *α*_1_ = *π/*4 ≈ 0.78, *α*_2_ = *π/*4 ≈ 0.78 and *K*_1_ = 0.08.

Further, we investigated the impact of the topology and network size on this emerging trend. Figs. 3(b) and 3(c) demonstrate that with increasing network size, the threshold value of *K*_2_ corresponding to the transition to asynchronous dynamics increase. Nevertheless, it is inevitable. Both these trends, i.e., the transition of network dynamics from phase synchrony to asynchrony corresponding to a threshold value of *K*_2_ and the increase in this threshold value with increasing network size is mirrored by random networks, as depicted in Figs. 3(d), 3(e) and 3(f). It is non-trivial that random networks undergo this transition as it implies that these trends are robust to network structure and network size. To solidify this claim, we plot the critical threshold value of *K*_2_ required for the system to transition to asynchronous dynamics against increasing network sizes in Fig. 4. Figure. 4(b) demonstrates that in case of random networks, even though the threshold *K*_2_ value might vary even for a fixed network size, it is due to randomness in the network and the trend of increasing in threshold values carry on nevertheless.

**FIG. 4.**
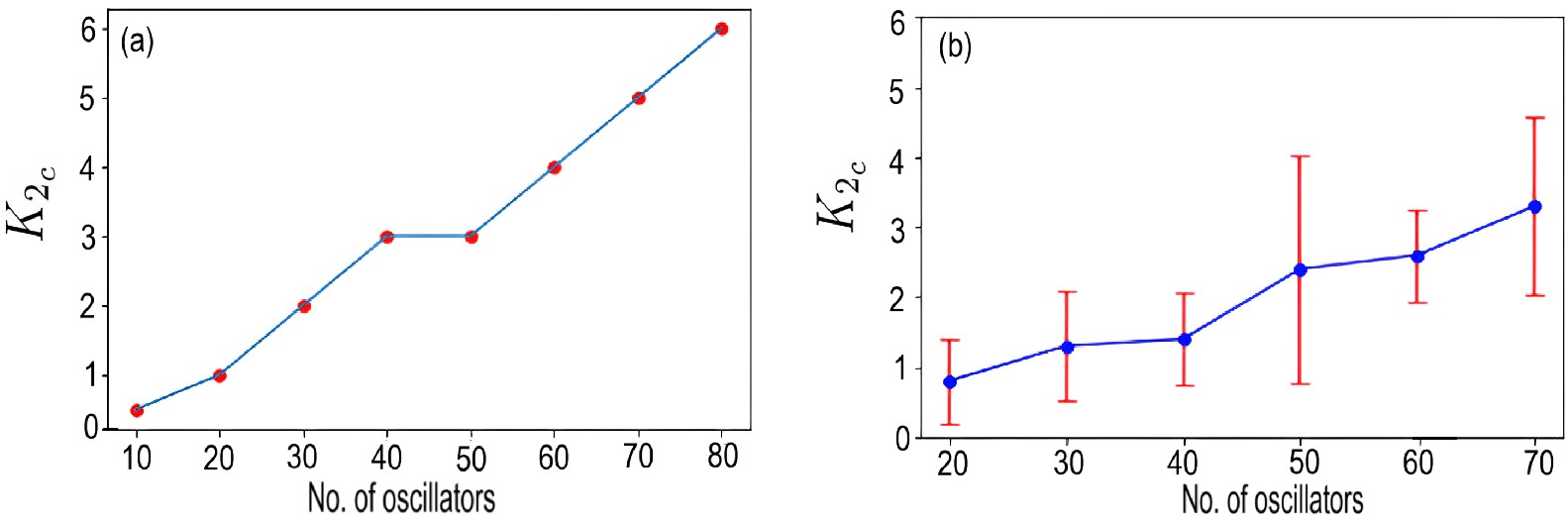
The threshold higher order coupling (*K*_2_) required for network to exhibit asynchronous dynamics plotted against increasing network sizes, once for (a) global network and once for (b) random network. The parameters are same as the previous figure.

Since very high asymmetry and higher-order coupling above a threshold can both induce asynchrony in the network dynamics, we investigate which one is the more dominant factor. Figure. 5(a) demonstrates the order parameter *Z* with parallel increase in first-order phase asymmetry *α*_1_ and higher-order phase aysmmetry *α*_2_ for a network of 20 nodes in absence of higher-order coupling. Evidently, only very large values of first-order asymmetry (*α*_1_) can induce asynchrony in network dynamics. It is unlikely that real ecosystems naturally and inherently favor a narrow range of very high asymmetry, irrespective of other conditions. However, Figs. 5(b) and 5(c) demonstrate that the interplay of increasing *K*_2_, very high *α*_1_ and high *α*_2_ induces asynchrony in the system. For comparable *K*1 and *K*2 values, asynchrony starts increasing. Increasing *K*_2_ values even further, there is a transition to complete asynchrony for high *α*_1_ and *α*_2_ as evidenced in Fig. 5(d). For intermediary values of *α*_1_ and *α*_2_, the degree of synchrony is also low. For low *α*_1_ and *α*_2_, very limited partial synchrony exists. For large *K*_2_, almost for the entire range of *α*_1_ and *α*_2_ the phase synchrony in network dynamics is extremely limited. In such a scenario, even for low *α*_1_ and *α*_2_, order parameter *Z* value is slightly more half of *Z* value corresponding to perfect synchrony. As shown in Fig. 5(e), when *K*_2_ values are increased even further, the level of partial phase-synchrony decreases even further. In effect, the entire dynamics is indeed asynchronous. Hence, the presence of higher-order interactions is vital and necessary for asynchronyous dynamics to exist.

**FIG. 5.**
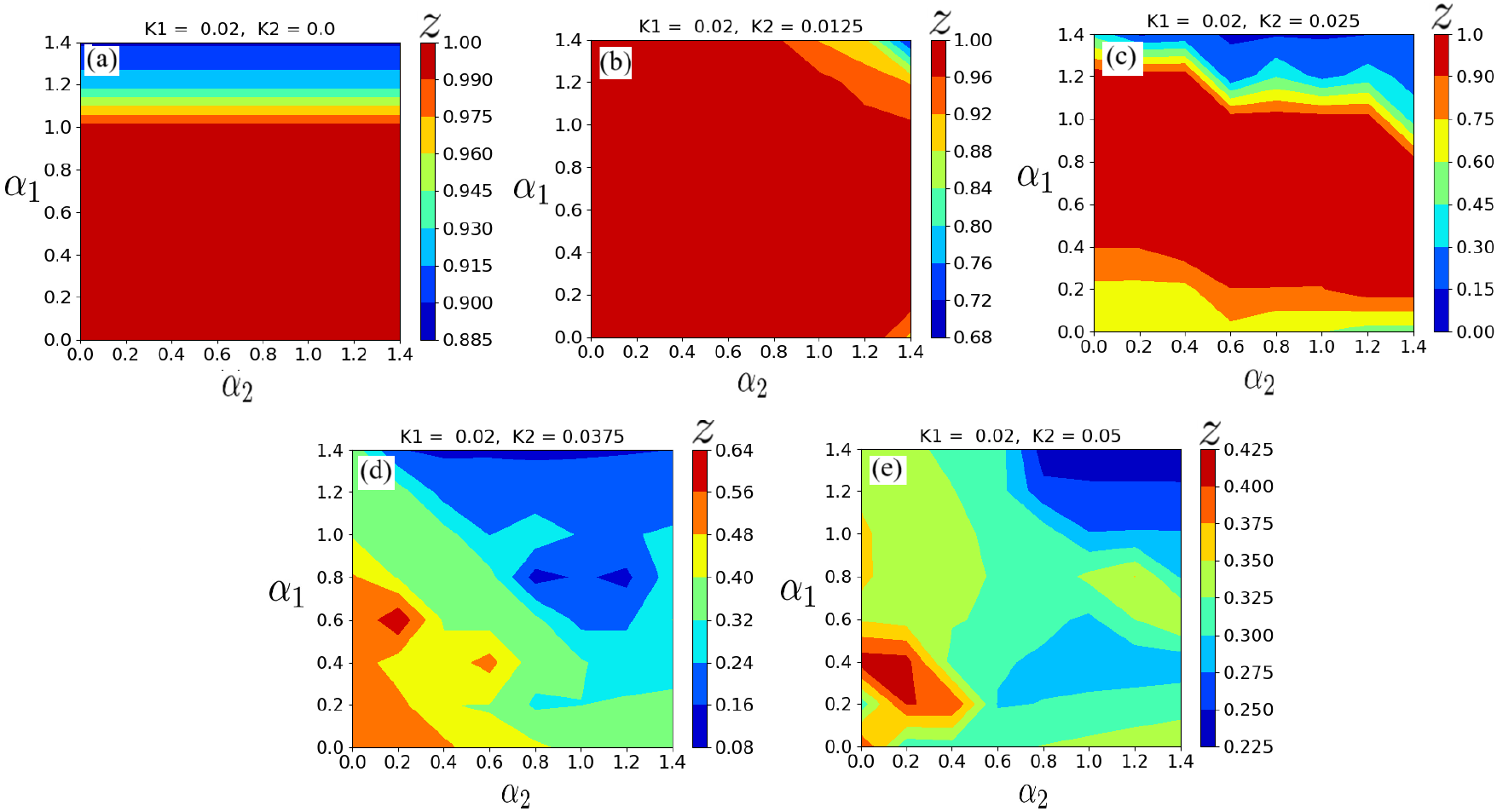
The order parameter variation with parallel increase of first-order phase assymetry *α*_1_ and higher-order phase asymmetry *α*_2_ for fixed first-order coupling strength *K*_1_ and higher-order coupling strength *K*_2_. The *K*_2_ value increases from (a)-(e).

### C. Impact of negative interactions on phase synchrony of ecological networks

As highlighted in the introduction, the inter-species interactions that occur naturally in an ecosystem may be non-trophic. Such interactions could be mutualistic or antagonistic and can be modelled as positive or negative couplings. Mathematically, these interactions can be represented respectively by positive and negative values of *K*_1_ and *K*_2_. So far, we have only considered positive coupling values. In this section, we analyze the role of negative coupling values on inducing asynchronous dynamics in the system.

#### 1. Impact of increasing percentage of negative first-order coupling

First, we measure the effect of negative first-order interactions on the ecosystem. Negative first-order coupling, represented by negative values of *K*_1_. We plot the order parameter of a 20 node network against increasing. First, we introduce 25% negative first-order couplings. As demonstrated in fig. 6(a), the oscillators formed two clusters for 75 percent positive values and 25 percent negative values, and the average order parameter value was lower than that in the case of fig.1(a). This implies that the network is partially phase synchronized. Figure. 6(b) shows that the system becomes completely asynchronous even for low values of *K*_1_ when 50% *K*_1_s are negative. On increasing the negative percentage of *K*_1_s further, i.e., 75% negative *K*_1_, we find that the oscillators are more asynchronously spread out. Figure. 6(c) demonstrates that presence of more positive first-order couplings than negative ones increase phase synchrony of network with increasing coupling amplitude, while the increase in percentage of negative first-order couplings leads to asynchronous dynamics in the network.

**FIG. 6.**
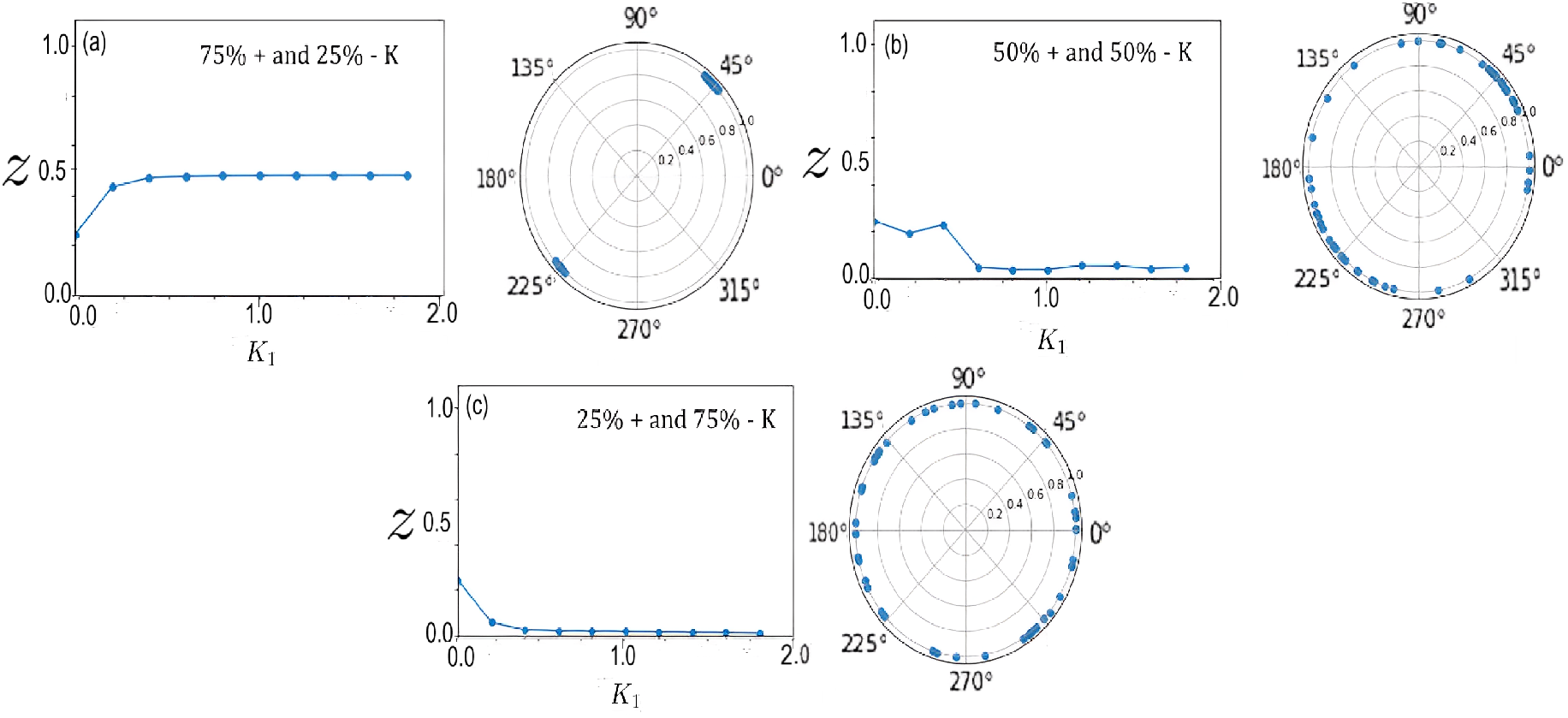
Order parameter *Z* vs first-order coupling (*K*_1_) for a global network of 20 nodes, with percentage of negative *K*_!_ coupling increasing from (a)-(c). The parameter values are *α*_1_ = *π/*4 ≈ 0.78, *K*_2_ = 0.

#### 2. Impact of increasing degree of negative higher-order coupling

Next, we analyze the effect of negative higher-order interactions (*K*_2_) on the ecosystem. To unerstand the behavior of the entire network, we plot the order parameter (*Z*) of the network corresponding to increasing percentage of negative coupling. We then repeat this exercise for gradually higher values of *K*_2_. Figure. 7 shows that over a particular threshold value of *K*_2_, there are specific ranges of *K*_2_ for which both high and low percentages of negative coupling that induce asynchrony in the system. This implies that both positive and negative *K*_2_ values induce asynchrony in the system, provided the interaction strength is over a threshold value. This result cements the role of higher order interactions in increasing the persistence of species-rich ecosystems.

**FIG. 7.**
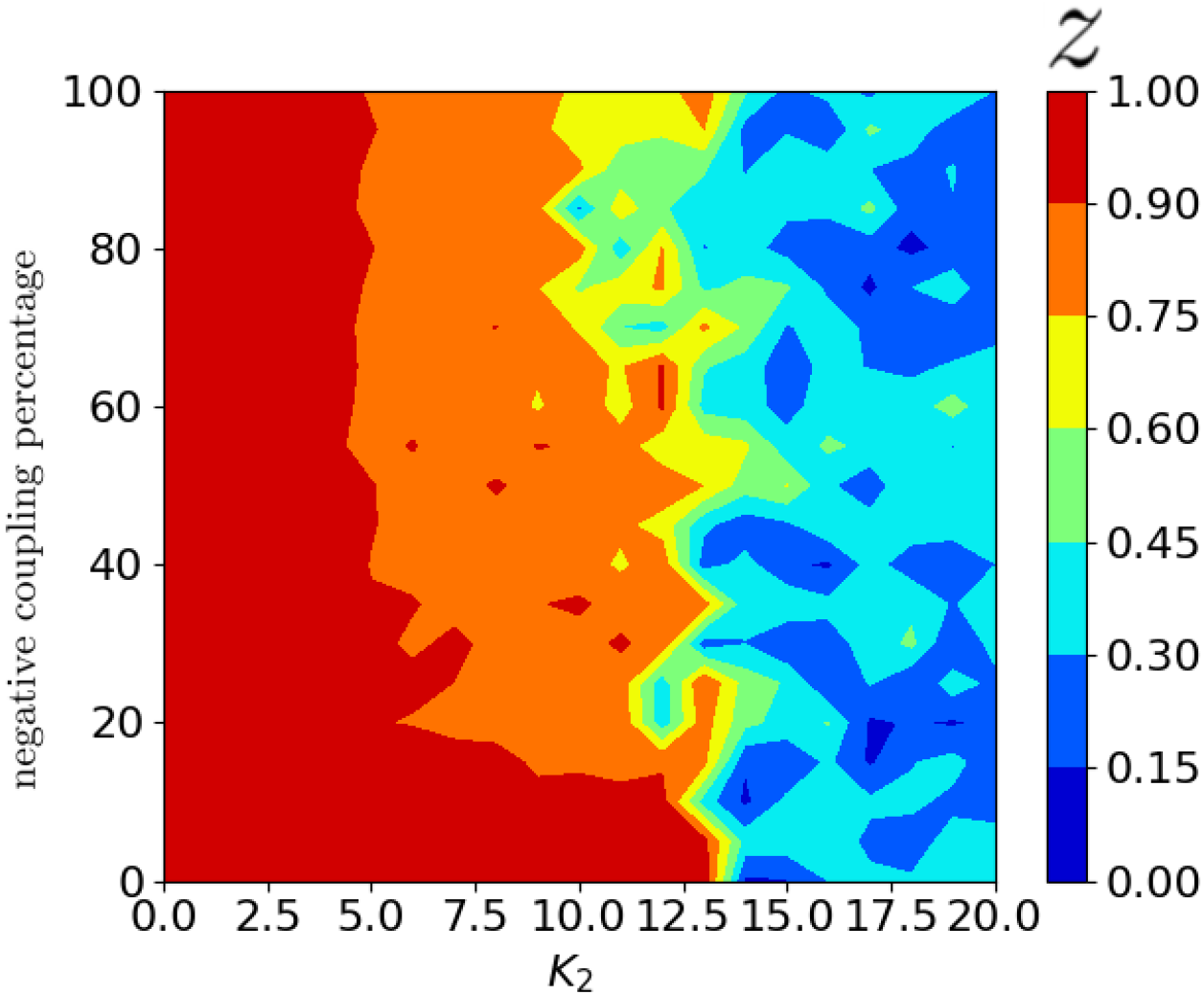
Order parameter *Z* vs increasing percentage of higher-order negative coupling *K*_2_ for a global network of 20 nodes. The pattern is repeated for increasing interaction strength of *K*_2_. The parameter values are *α*_1_ = *π/*4 ≈ 0.78, *α*_2_ = *π/*4 ≈ 0.78 and *K*_1_ = 0.04.

#### 3. Impact of increasing degree of negative first-order and higher-order coupling

Clearly, both negative first-order coupling (*K*_1_) and negative higher-order coupling (*K*_2_) seperately play a role in making the dynamics of an entire network asynchronous. However, based on our discussion, it is vital to understand how negative couplings in both first (− *K*_1_) and higher-order (− *K*_2_) simultaneously impact the dynamics of the network. To analyze this, we plot a heatmap of the order parameter *Z* of the network with simultaneously increasing values of *K*_1_ and *K*_2_, as shown in Fig. 8. The percentage of both negative first-order and negative higher-order couplings increase from Figs. 8(a)-(f). Figure 8(a) demonstrates that for no negative coupling (i.e., negative coupling % = 0), asynchrony is induced in the network only when *K*_2_ is larger than *K*_1_. Next, Fig. 8(b) demonstrates that with slight increase in negative coupling percentage of both *K*_1_ and *K*_2_ (i.e., negative coupling % = 5), asynchrony is induced in the system at a lesser threshold value of *K*_2_. On increasing the negative coupling percanetage of both *K*_1_ and *K*_2_ to 25% (refer Fig. 8(c)), it is observed that not only does the red region signifying the phase synchronized dynamics of network decreases vastly. Equally importantly, there is no scope of perfect phase synchrony. Indeed, the highest possible value that the order parameter *Z* attains is 0.7, depicting only partial phase synchrony. When we increase the percentage of negative *K*_1_s and *K*_2_s to 50, the region of high, though partial phase-synchrony gets decreased further, and the degree of synchrony is even lesser than the previous case as illustrated in fig.8(d). Typically, when majority of the *K*_1_ and *K*_2_ couplings are negative (75%), the degree of partial phase-synchrony decreases further. Barring a very few randomly distributed pockets of partial phase-synchrony, for both high and medium *K*_2_ and high *K*_1_, the network exhibits asynchronous dynamics as demonstrated by fig.8(e).

**FIG. 8.**
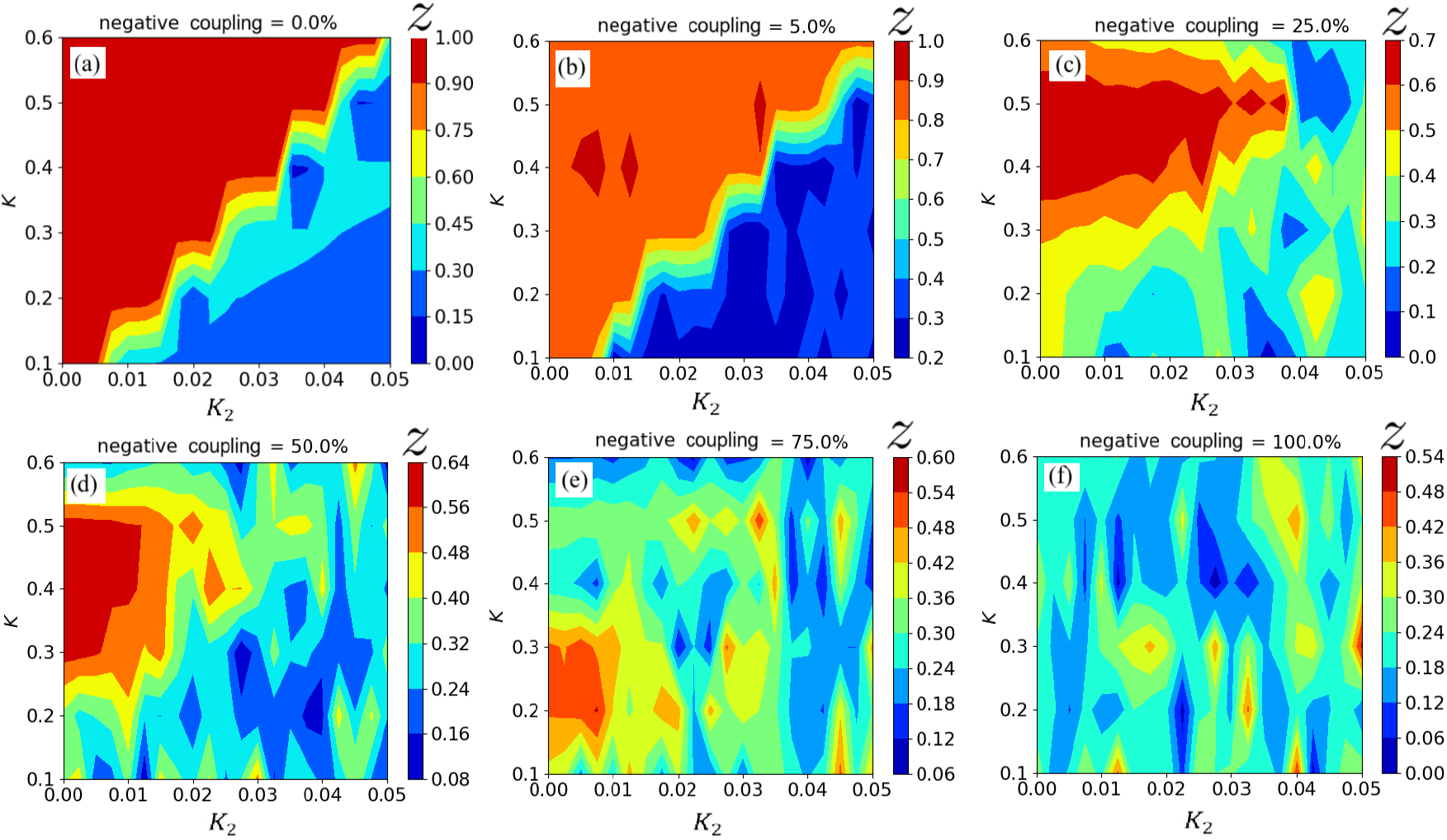
Order parameter vs simulatenous increase in first-order coupling (*K*_1_) and higher-order coupling (*K*_2_) for a global network of 20 nodes with (a) 0% negative coupling in both *K*_1_ and *K*_2_, (b) 5% negative coupling in both *K*_1_ and *K*_2_, (c) 25% negative coupling in both *K*_1_ and *K*_2_, (d) 50% negative coupling in both *K*_1_ and *K*_2_, (e) 75% negative coupling in both *K*_1_ and *K*_2_ and (f) 100% negative coupling in both *K*_1_ and *K*_2_. The parameter values are *α*_1_ = *π/*4 ≈ 0.78, *α*_2_ = *π/*4 ≈ 0.78.

## IV. DISCUSSION AND CONCLUSION

Species-rich ecosystems consist of numerous interacting species living in close proximity. Therefore, an ecosystem must be accurately modelled as a large complex network, with each species acting as nodes of the network and the inter-species interactions acting as its links. Most of the ecological inter-species interactions are non-linear, with the non-linearity inherent to the modelling approach on which the network is based. Specifically, prey-predator models incorporate such inherent non-linearity within the equations which describe the population dynamics. Population dynamics of a variety of ecosystems modelled as prey-predator models exhibit an oscillatory behavior. If such oscillations are phase synchronized, the persistence of the ecosystem is lowered as the total population reaches low density values simultaneously and is therefore more susceptible to external factors like environmental noise, Moran effect (environmental conditions) or disease outbreaks. This could lead to the entire population going extinct through cascading effect. In contrast, if the population dynamics is asynchronous, that implies increased persistence due to increased chance of community survival through rescue effect. Therefore, to measure extinction risk of an ecosystem, an investigation measuring the total phase dynamics of all the constituent population is extremely insightful. Previous studies on paradigmatic prey-predator models establish that the phase dynamics of a prey-predator pair can be accurately approximated using a phase oscillator model. Thus, an ecological network can be parallely modelled as a network of phase oscillators, linked to each other through interactions of various strengths.

However, constructing such a network requires specific considerations. The close proximity of a multitude of species in the ecological network ensures that interspecies interactions are not exclusively pairwise. To account for numerous likely interactions involving more than two species, higher-order interaction terms need to be included in the equation of each phase oscillator. Further, to account for inherent habitat heterogeneity of an ecosystem, the includsion of asymmetrical phase in the dynamics of the oscillators is also very important. Motivated by the above, we model an ecosystem as a a speciesrich ecological network, where each node consists of a Sakaguchi-Kuramoto phase oscillator model. We investigate the phase dynamics and degree of phase synchronization between the nodes of this network and commment on the persistence of this network.

Our studies reveal that even with acceptable levels of asymmetry in the dynamics of each node, the network still exhibits strong phase synchrony under high first-order coupling. Only very high first-order asymmetry phase can induce limited asynchrony in absence of higher-order interactions. However,we find that introducing the higher-order interaction term changes the network dynamics drastically. Higher-order coupling over a threshold amplitude makes the network dynamics asynchronous. With increasing network size, this degree of higher-order coupling required to induce asynchronicity in the network increases. We observe this trend for both globally connected and random networks, proving that this trend is robust to network structure. Next, we incorporated various mutualistic (+) and antagonistic (-) inter-species interactions through positive and negative coupling values in both first and higher-order. We observe that in a network of fixed size, if both first and higher order coupling have a combination of positive and negative values, asynchrony of the network is strongly dependent on both the percentage of first and higher-order negative couplings. We find that with increasing percentage of negative coupling, phase-asynchrony in the network is promoted.

The continued existence of ecosystems strongly suggest the presence of asynchrony in the dyanmics of each species within ecological networks. A primary future goal of this research is to analyze ecological networks of increasing complexity with this approach. For example, environmental noises can be introduced in the dynamics of each phase oscillator and its effect on the persistence of ecosystems can be investigated. The same could be done by incorporating further higher-order interactions and/or considering multi-layer networks. Analysis of the synchrony groups and chimeric elements that emerge from the model could provide insights into the empirical factors contributing to the formation of these groups. The relationship between chimeric elements and subgroups formed by the nodes, and its impact on persistence of the ecological network similarly merit an investigation.

